# Examination of rickettsial host range for shuttle vectors based on *dnaA* and *parA* genes from the pRM plasmid of *Rickettsia monacensis*

**DOI:** 10.1101/2022.02.09.479837

**Authors:** Nicole Y. Burkhardt, Lisa D. Price, Xin-Ru Wang, Chan C. Heu, Gerald D. Baldridge, Ulrike G. Munderloh, Timothy J. Kurtti

**Affiliations:** University of Minnesota, Department of Entomology, Saint Paul, Minnesota, USA

**Keywords:** *Rickettsia* plasmids, *Rickettsia monacensis*, *Rickettsia amblyommatis*, Shuttle plasmids, Transformation, Host range

## Abstract

The genus *Rickettsia* encompasses a diverse group of obligate intracellular bacteria that are highly virulent disease agents of mankind as well as symbionts of arthropods. Native plasmids of *Rickettsia amblyommatis* (AaR/SC) have been used as models to construct shuttle vectors for genetic manipulation of several *Rickettsia* species. Here we report on the isolation of the complete plasmid (pRM658B) from *Rickettsia monacensis* (IrR/Munich) mutant Rmona658B and the construction of shuttle vectors based on pRM. To identify regions essential for replication, we made vectors containing the *dnaA* and *parA* genes of pRM with varying portions of the region surrounding these genes and a selection-reporter cassette conferring resistance to spectinomycin and expression of green fluorescent protein. *Rickettsia amblyommatis* (AaR/SC), *R. monacensis* (IrR/Munich), *Rickettsia bellii* (RML 369-C), *Rickettsia parkeri* (Tate’s Hell), and *Rickettsia montanensis* (M5/6) were successfully transformed with shuttle vectors containing pRM *parA* and *dnaA*. PCR assays targeting pRM regions not included in the vectors revealed that native pRM was retained in *R. monacensis* transformants. Determination of native pRM copy number using a plasmid-encoded gene (RM_p5) in comparison to chromosomal encoded *gltA* indicated reduced copy numbers in *R. monacensis* transformants. In transformed *R. monacensis*, native pRM and shuttle vectors with homologous *parA* and *dnaA* formed native plasmid-shuttle vector complexes. These studies provide insight on the maintenance of plasmids and shuttle vectors in rickettsiae.

**IMPORTANCE:**

- This paper describes a new series of plasmid shuttle vectors for the transformation of rickettsiae.
- Shuttle vectors based on *parA* and *dnaA* sequences of the plasmid pRM can be used to transform diverse rickettsia as well as its native host *Rickettsia monacensis*.
- Our results provide insight into the maintenance of rickettsial-based shuttle vectors in rickettsiae.

---

The genus *Rickettsia* (Order *Rickettsiales*, Family *Rickettsiaceae*) contains obligate intracellular α-proteobacteria with wide ranging effects on infected invertebrates and mammals: some such as *Rickettsia tamurae* subspecies *buchneri* are avirulent arthropod symbionts while others (for example, *R. rickettsii* and *R. prowazekii*) are highly virulent vector-borne human pathogens causing spotted fevers and typhus. Some *Rickettsia* spp. are unique among the *Rickettsiales* in harboring plasmids (1). More than 30 plasmids have been identified in a wide range of *Rickettsia* species and strains (2 - 6). The diversity of *parA* on rickettsial plasmids suggests that foreign plasmids have invaded rickettsiae (7) via a conjugation system (8, 9). As many as 4 distinct plasmids, each with a distinct partitioning system, can coexist in a given rickettsial strain, (2, 10). Plasmids are more commonly associated with avirulent and less virulent rickettsiae and are absent in highly virulent strains (Table S1). Rickettsial plasmids have undergone reductive evolution and to date, virulence has not been linked to presence of plasmids (5, 11). These features indicate that shuttle vectors modeled after rickettsial plasmids can be developed as tools to introduce foreign genes into rickettsiae, as we have previously demonstrated previously (12).

The discovery of low-copy number plasmids in many *Rickettsia* spp. (1, 2, 13) spurred development of the first shuttle vectors, a fundamental advance in rickettsial transformation technologies (12, 14). Their potential has been realized by generation of transformant rickettsiae expressing selectable markers and fluorescent proteins that have enabled study of rickettsial interactions with host cells in unprecedented detail. Examples with arthropod endosymbionts include *R. t. buchneri* and *R. peacockii* in tick host cells (15), as well as *R. bellii* expressing a plasmid-encoded heterologous *rick*A gene (16). Although the typhus group are among those *Rickettsia* spp. that do not harbor a native plasmid, shuttle vectors have also been used to transform and study typhus group rickettsiae, including the epidemic typhus agent, *R. prowazekii*, in L929 murine fibroblast cells (17) and the endemic typhus agent, *R. typhi*, in CB17 SCID mice and in association with CD8 cells (18). The first generation rickettsial shuttle vectors are now an important component of the still limited toolbox for genetic manipulation of rickettsiae (19, 20), underscoring the need for further expansion and optimization of the vectors, and for the study of plasmid biology in the genus *Rickettsia*.

The design of plasmid-based bacterial transformation vectors was influenced by discovery of incompatibility, which resulted in classification of plasmids into incompatibility groups determined by their ability to be simultaneously maintained within a single host cell. Incompatibility arises when both foreign and native plasmids carry the same or a similar *parA* gene, causing both plasmids to become unstable (21, 22). *R. t. buchneri* and *R. amblyommatis*, have multiple independent plasmids, each with a distinct *parA* (2, 3, 10). The role of *parA* in plasmid partitioning and incompatibility in rickettsiae remains largely unexplored and has likely practical consequences for use of rickettsial shuttle vectors. In contrast, *R. monacensis* contains only one native plasmid, pRM, and phylogenetic analysis shows it contains a *parA* with no significant similarity to those from 19 other rickettsial plasmids (3), although the pRAS01 from *R. asembonensis* strain NMRCii has a *parA* with 66% nucleotide identity to *parA* from pRM (23). The predicted amino acid sequence of ParA from pRM is most similar to ParA-like predicted proteins from *Mycobacterium* and *Bartonella* spp. (25-53% BLASTP identities). Thus, shuttle vectors from pRM should theoretically be compatible for transformation of an expanded range of rickettsiae.

Our aim is to both expand the repertoire of shuttle vectors for use in the study of *Rickettsia* spp. and improve our understanding of the mechanisms of plasmid maintenance in diverse rickettsial species. Here we report isolation of pRM from *R. monacensis* in *Escherichia coli* and its development as a family of shuttle vectors, which were used to transform several rickettsiae including *R. monacensis* and *R. amblyommatis* AaR/SC, whose pRAM plasmids have also been developed as rickettsial shuttle vectors (12). Quantitative PCR results indicated that introduction of the pRM shuttle vector reduced copy numbers but did not eliminate endogenous pRM in transformed *R. monacensis*. The first-generation pRAM- and second-generation pRM-based shuttle vector families carry unrelated *parA* genes and thus diversify the range of vector choices available for transformation of rickettsiae.

## RESULTS

### Identification of minimal coding sequences required for pRM shuttle vector replication and partitioning

Because coding sequences that support plasmid replication and partitioning are often clustered together, we identified the minimal region required for rickettsial shuttle vector replication and partitioning by constructing three shuttle vectors with varying portions of RM_p16-21 coding sequences: pRMdSGK Clone #1 [pRMΔ1], pRMdSGK Clone #2 [pRMΔ2] and pRMdSGK Clone #3 [pRMΔ3] (Fig.1). All three clones contained coding sequence for the pRM DnaA-like protein (RM_p16) and ParA (RM_p18) that function in DNA replication and chromosomal stability, as well as RM_p17 and RM_p19, hypothetical proteins (HP) of unknown function, although RM_p19 contains domains with similarity to the HTH_XRE family transcriptional regulators. The smallest test construct, pRMΔ1, included the gene cluster RM_p16 through RM_p19 and partial coding sequence (359 bp of 507 bp) for RM_p20 (green bar Fig. 1C), a second likely HTH_XRE family protein with 74% similarity to RM_p19. The pRMΔ2 construct extended the same *dnaA*/*parA* region through an intact RM_p20 coding sequence and an amino terminal fragment of RM_p21 (orange bar Fig. 1C). The pRMΔ3 construct further encoded an intact RM_p21, a likely Sca12 cell surface antigen, and approximately 1 kbp of downstream non-coding sequence (lilac bar Fig. 1C).

**FIG 1.**
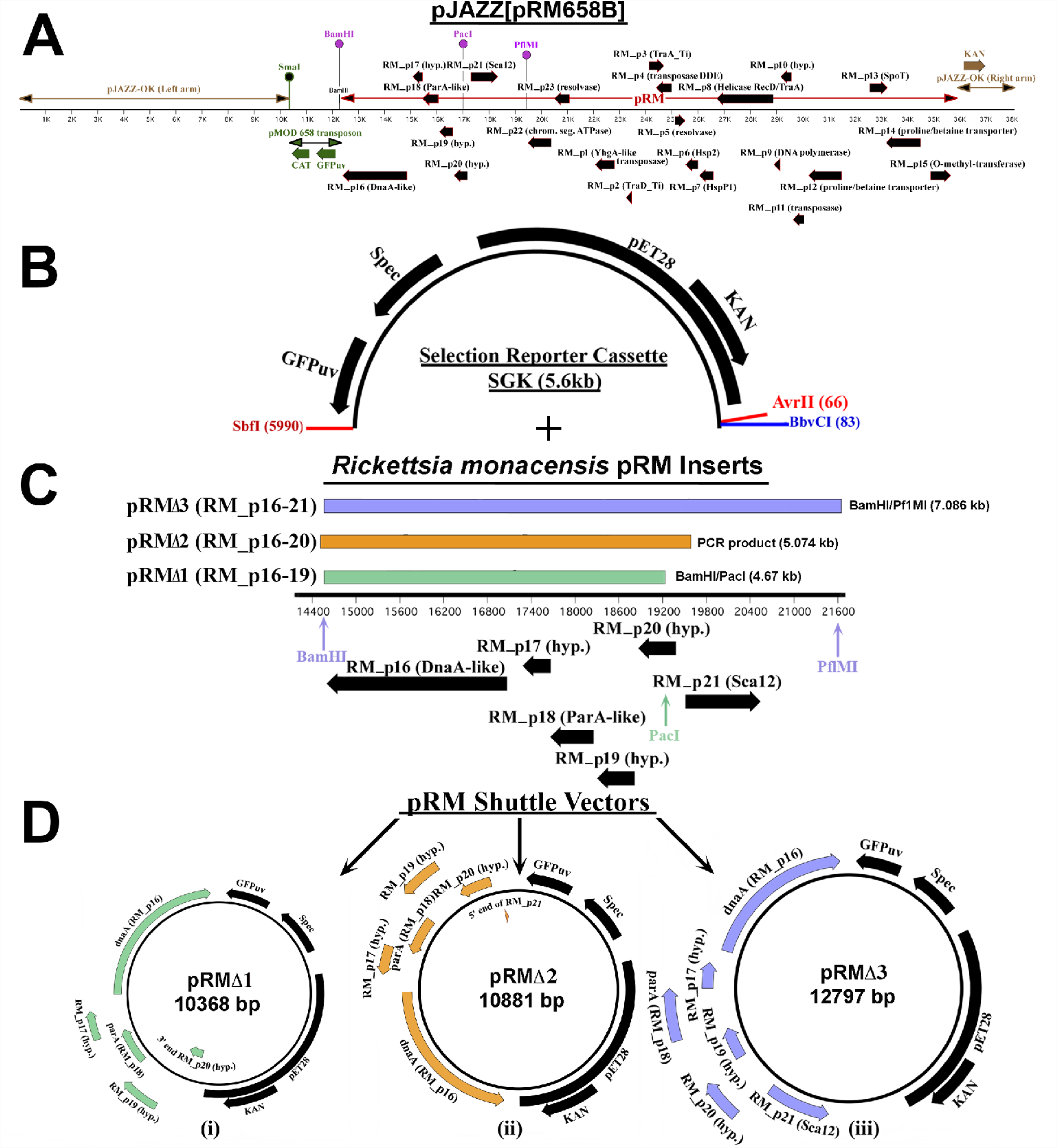
Construction of *R. monacensis* pRM shuttle vectors. (A) Schematic diagram depicting construct pJAZZ[pRM658B], the *R. monacensis* plasmid pRM with pMOD658 transposon cloned into the linear plasmid pJAZZ. The two arms formed by digested pJAZZ are indicated by brown double-ended arrows and pRM is represented by a red double-ended arrow; genes present on pRM are denoted with black arrows. The pMOD658 transposon is shown in green; resistance to chloramphenicol (CAT) and the ability of the clone to express green fluorescent protein (GFP_uv_) are conferred by the transposon inserted in pRM. The unique SmaI site with which pRM was linearized for cloning is located in the transposon. Pink balloons indicate the restriction enzyme sites used for sub-cloning the *dnaA*/*parA* region of pRM. (B)The 5.6 kbp selection reporter cassette SGK was ligated with three pRM fragments of varying size containing *dnaA* and *parA* yielding the *R. monacensis* pRM shuttle vectors pRMΔ1, 2 and 3 (C). The numbered black line in panel C represents bp 14400 to 21600 of the 23486 bp *R. monacensis* plasmid pRM and the thick black horizontal arrows beneath it indicate predicted genes and their orientations. Colored vertical arrows indicate the restriction enzyme sites used during pRM sub-cloning and colored bars above the numbered black line represent the three fragments of pRM contained in the finished shuttle vectors shown in (D).

### Transformations trials of five SFG *Rickettsia* spp

The smallest test construct, pRMΔ1, transformed only *R. monacensis* (Table 1), which carries pRM as its native plasmid. In contrast, trials with pRMΔ2 (encoding the same RM_p16 through RM_p19 proteins as well as the intact RM_p20 XRE family protein with its upstream region) and pRMΔ3 (extended to encode the RM_p21 Sca12 cell surface antigen), transformed all five *Rickettsia* species tested (Table 1). Species successfully transformed included four SFG rickettsiae, notably *R. amblyommatis* AaR/SC which contains 3 native plasmids, as well as *R. bellii*, which occupies a more basal phylogenetic position (24). The RM_p16 through RM_p19 cluster thus supported transformation of the parental *R. monacensis*, but inclusion of the RM_p20 locus and/or a short upstream sequence extended the range of a pRM-based shuttle vector to other SFG and ancestral group rickettsiae. In contrast to RM_p20, RM_p21 and the associated non-coding 1 kbp sequence conferred no apparent advantage.

**TABLE 1.** Transformation of *Rickettsia* with plasmid constructs containing pRM *dnaA* and *parA*

| CONSTRUCT |  |  |  |  |
| --- | --- | --- | --- | --- |
| Code | pRAM18dRGA | pRMΔ1 | pRMΔ2 | pRMΔ3 |
| pRM genes included | --- | (RM_p16-20<sup>1</sup>) | (RM_p16-21) | (RM_p16-21) |
| Size | (10.3 kbp) | (10.4 kbp) | (10.9 kbp) | (12.8 kbp) |
| **<u>STRAINS WITH ENDOGENOUS PLASMIDS</u>** |  |  |  |  |
| *R. monacensis* (IrR/Munich) | + | +<sup>2</sup> | + | + |
| (pRM<sup>3</sup>, 23<sup>4</sup>) |  |  |  |  |
| *R. amblyommatis* (AaR/SC) | + | -<sup>5</sup> | + | + |
| (pRAM<sup>3</sup>, 18<sup>4</sup>, 23<sup>4</sup>, 32<sup>4</sup>) |  |  |  |  |
| **<u>PLASMID FREE STRAINS</u>** |  |  |  |  |
| *R. parkeri* (Oktibbeha) | + | - | + | + |
| *R. montanensis* (M5/6) | + | - | + | + |
| *R. bellii* (369C) | + | - | + | + |

<sup>1</sup> Italicized gene number indicates sequence does not encode complete gene
<sup>2</sup> Transformed
<sup>3</sup> Native plasmid name
<sup>4</sup> Approximate native plasmid size (kb)
<sup>5</sup> Not transformed

These results indicated that the RM_p16 through RM_p20 loci of pRM contained the minimum necessary DNA sequence and protein coding capacities for a shuttle vector capable of transforming a wide range of SFG and ancestral group rickettsiae. The promoter-prediction program BPROM (25) indicated a single promoter upstream of the RM_p20 locus but none for RM_p17, 18 or 19, consistent with functional dependence of RM_p18 expression on sequences upstream of RM_p20 and a likely operon extending through the RM_p16 locus (Fig 1D). The predicted transcription start site was at bp 19395 of pRM, with a -10 box from 19402-19410, and a -35 box from 19425-19430. A comparison of growth rates of wild-type (WT) *R. monacensis* and pRMΔ2-transformed *R. monacensis* using three separate growth curve analyses indicated no significant differences between the growth rates of WT and transformants (doubling times of 17.5 and 18 hours, respectively).

### Evaluation of GFP expression in wild-type and transformed *R. monacensis* using confocal microscopy

Cell free *R. monacensis* WT and transformants were stained with NucBlue^™^ Live Cell Stain ReadyProbes^™^ (Thermo Fisher Scientific, Waltham, MA) to visualize all rickettsiae whether transformed or not, then gently mounted onto slides using a Cytospin centrifuge (Thermo Fisher) and observed using confocal microscopy. Column 1 of Figure 2A with a DAPI filter shows all the rickettsiae present, while column 2 shows only *gfp*_*uv*_-expressing transformed rickettsiae (FITC filter). Column 3 illustrates the overlap of fluorescence in the DAPI and FITC fields. Visually, the degree of co-localization indicated that nearly all the rickettsiae present were transformed. To confirm these observations, we used Pearson’s coefficient (PCC) and Manders’ co-localization coefficient (MCC) (26, 27) to evaluate the co-localization of the DAPI and FITC fluorescence. For the PCC analysis, the correlation coefficient was measured on all pixels in an individual image for three random fields. The PCC assay values range from -1 to 1 with 1 being 100% co-localization; the PCC values are lower than would be indicated by visual analysis as low fluorescence values limit the program’s ability to evaluate and interpret pixels. The MCC assay measures the proportion of rickettsial DNA fluorescence that localizes with transformed plasmid fluorescence (M1) and vice versa (M2). Almost 100% co-localization is indicated by the very similar M1 and M2 values calculated for each of the three transformed *R. monacensis* (Figure 2C).

**FIG 2.**
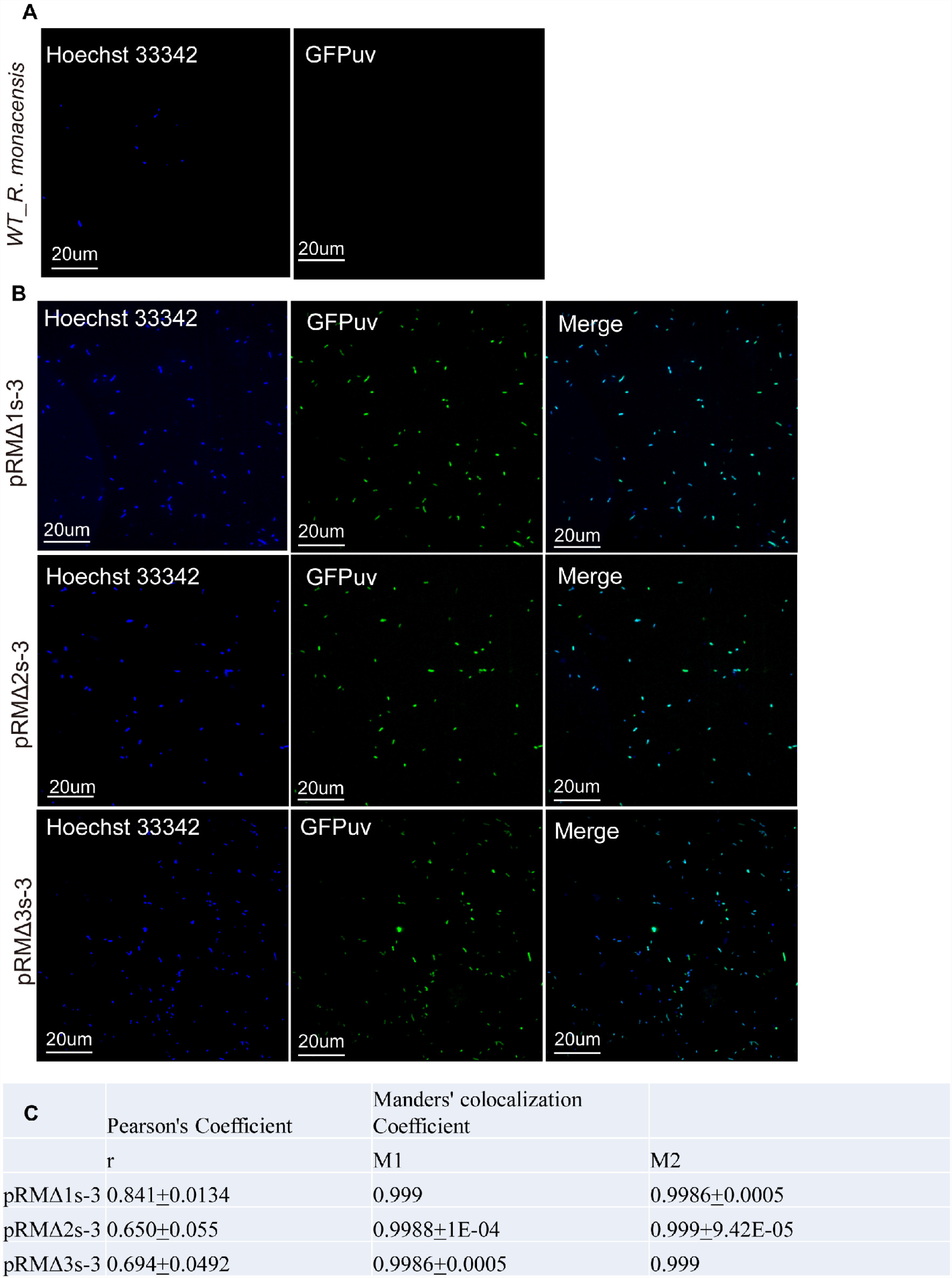
Evaluation of GFPuv expression in WT and transformed *R. monacensis* using confocal microscopy. (A) WT and (B) pRMΔ1, 2 and 3-transformed *R. monacensis*. The first column shows fluorescent staining of rickettsial DNA (blue signal, Hoechst 33342) and the second column shows *gfp*_*uv*_ fluorescence (green signal). The third column shows combined Hoechst 33342 and *gfp*_*uv*_ signals of the merged fields. (C) Tables showing Pearson’s correlation coefficient (PCC) and Manders’ colocalization coefficient (MCC) of same fields of view shown in (B) to quantify co-localization of *gfp*_*uv*_ fluorescence from transformed *R. monacensis* and fluorescence from stained rickettsial DNA. M1: fraction of blue fluorescence in areas with green fluorescence; M2: fraction of green fluorescence in areas with blue fluorescence.

### Presence of shuttle vector in pRM-transformed *R. amblyommatis* and *R. parkeri* and conservation of native plasmids in transformed *R. amblyommatis*

Undigested DNA from *R. amblyommatis* and *R. parkeri* transformed with pRMΔ2 and pRMΔ3 was separated by pulsed-field gel electrophoresis (PFGE) (Fig. 3A) and transferred onto zeta probe membranes. Because *R. parkeri* does not contain native plasmids, hybridization of *R. parkeri* pRMΔ2 and pRMΔ3 transformants with digoxigenin-labeled *gfp*_*uv*_ probe (28) identified the unaltered shuttle vectors with asterisks indicating the nicked linear forms of pRMΔ2 and pRMΔ3 at 10.81and 12.814 kbp, respectively (Fig.3B). The *R. amblyommatis* transformants show a band pattern similar to that of *R. parkeri*; however extra bands were present in both *R. amblyommatis* pRMΔ2 and pRMΔ3 hybridized with the *gfp*_*uv*_ probe (Fig. 3B). Stripped Southern blots were hybridized with recombinase, *hsp2* and helicase digoxigenin-labeled probes specific for *R. amblyommatis* strain AaR/SC plasmids pRAM18, pRAM23 and pRAM32, respectively (12). Bands of the appropriate size were observed for all 3 native pRAM plasmids (pRAM18 = 18.344 kbp; pRAM23 = 22.852 kbp and pRAM32 = 31.972 kbp) in the transformed *R. amblyommatis* (Fig. 3A, C, D and E; marked with asterisks), but not in *R. parkeri*. Thus, pRM shuttle vector and all three native *R. amblyommatis* plasmids are conserved in the pRM transformed *R. amblyommatis*.

**FIG 3.**
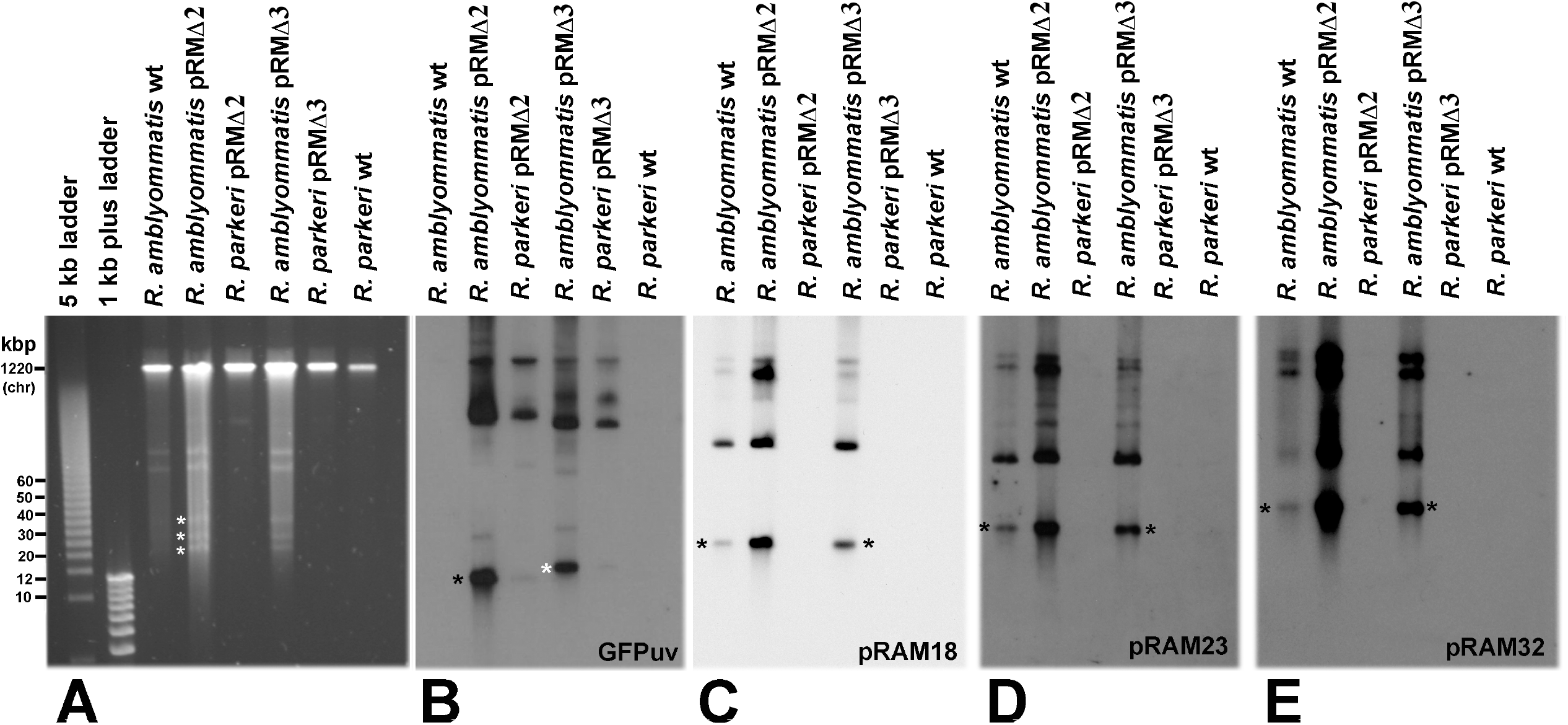
Pulsed-field gel electrophoresis and Southern blot analysis of *R. amblyommatis* and *R. parkeri* transformed with pRM shuttle vectors pRMΔ2 and pRMΔ3. *gfp*_*uv*_ and pRAM digoxigenin-labeled probes were used to detect the presence of shuttle vector, pRAM18, pRAM23 or pRAM32 in transformants. (A) PFGE gel. White asterisks mark the nicked linear forms of pRAM18, pRAM23 and pRAM32 respectively. (B) Southern blot analysis of pulsed-field gel hybridized with digoxigenin-labeled *gfp*_*uv*_ probe. The nicked linear forms of pRMΔ2 and pRMΔ3 are denoted by black and white asterisks, respectively. Higher molecular weight bands are supercoiled and multimer forms of the shuttle vectors pRMΔ2 and pRMΔ3. Note absence of labeling in the WT *R. amblyommatis* and *R. parkeri* lanes. (C) Southern blot analysis of panel A hybridized with digoxigenin-labeled pRAM18 probe (invertase). (D) Southern blot analysis of panel A hybridized with digoxigenin-labeled pRAM23 probe (Hsp2). (E) Southern blot analysis of panel A hybridized with digoxigenin-labeled pRAM32 probe (RecD). Black asterisks in panels C, D and E denote nicked linear forms of pRAM18, 23 and 32, respectively.

### Testing for the presence of native pRM and shuttle vectors in pRMΔ-transformed *R. monacensis*

To assess native pRM in shuttle vector-transformants, genomic DNA from WT and transformant *R. monacensis* was PCR-amplified with native pRM-specific primer sets (Table 2). All three *R. monacensis* transformants contained native pRM as indicated by the presence of amplicons from genes present in native pRM but not in the shuttle vectors (Fig. 4A, B & C). The dGFPuvF2/R2 primer amplicons confirmed presence of the shuttle vectors (Fig. 4D). *R. amblyommatis* AaR/SC pRMΔ2 DNA was used as a control to confirm primer specificity; as expected, it yielded no amplicon with native pRM primer sets but was positive with dGFPuvF2/R2 primers.

**TABLE 2.**
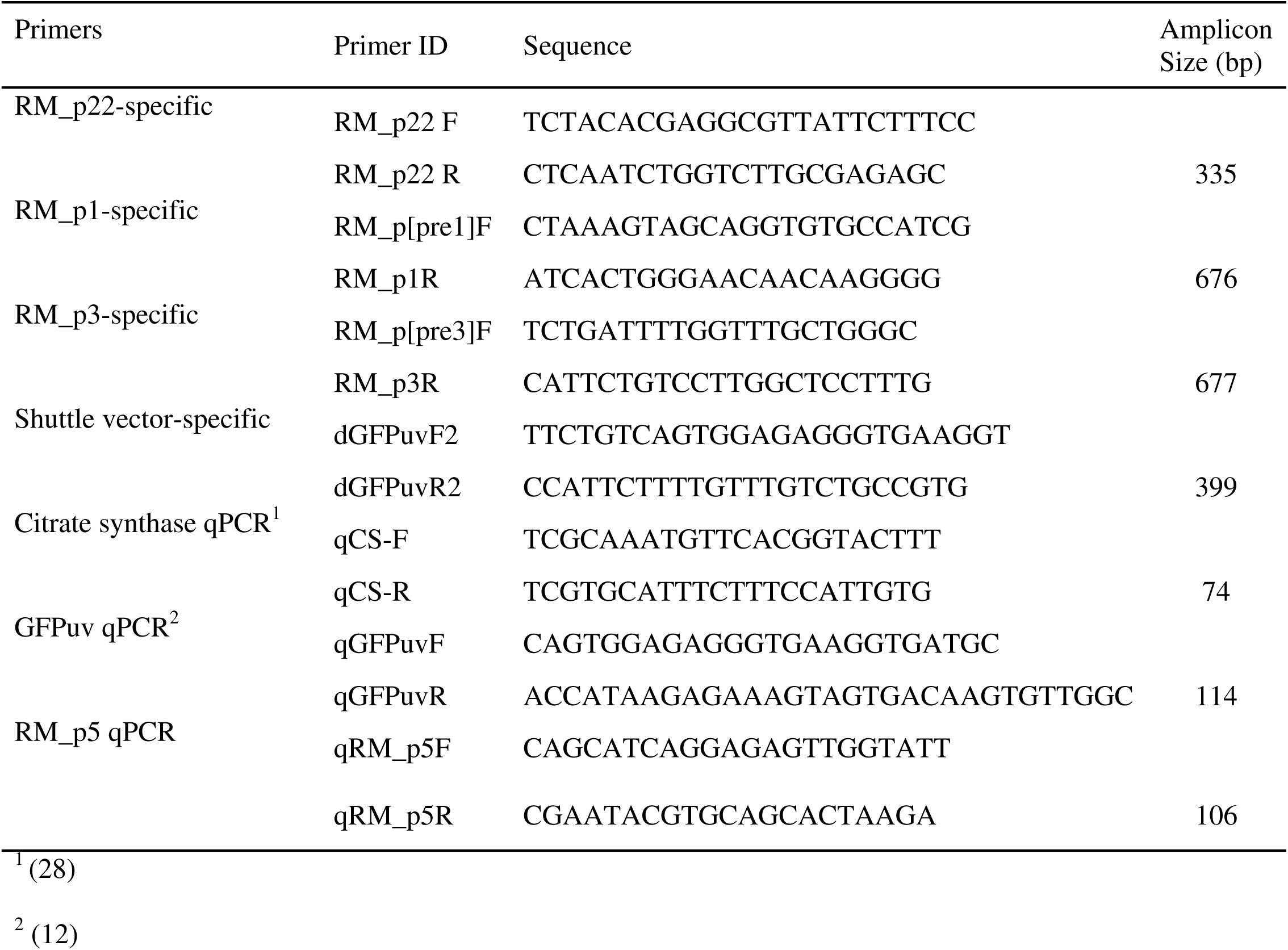
PCR and qPCR primers used to distinguish native pRM and shuttle vectors and estimate their copy numbers

**FIG 4.**
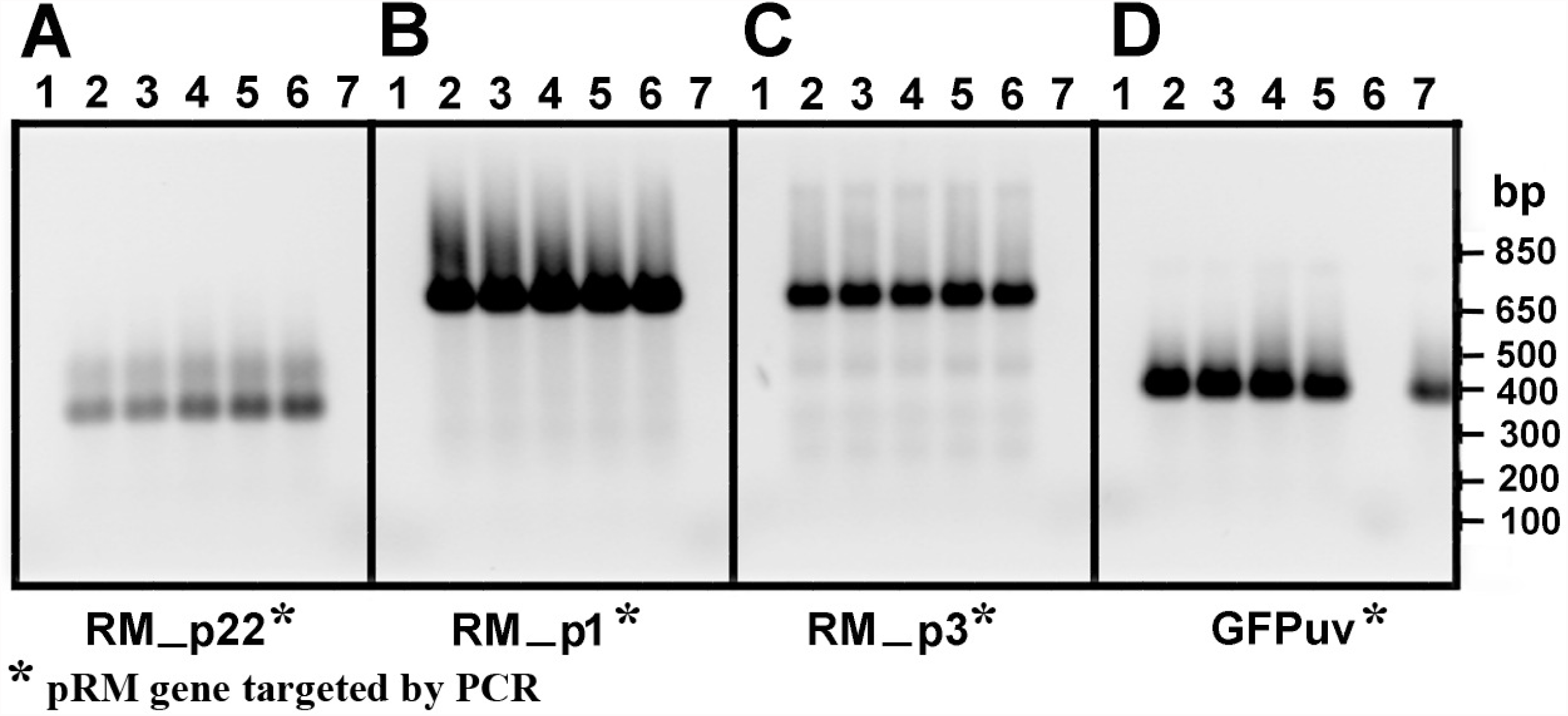
PCR results demonstrating the presence of native pRM in pRMΔ-transformed *R. monacensis*. PCR amplicons were electrophoresed on a 1% agarose gel stained with GelGreen (Biotium, Hayward CA). (A), (B), & (C) PCR results using primers specific for native pRM genes (RM_p22, RM_p1, and RM_p3) not present in pRM shuttle vectors. (D) PCR products using *gfp*_*uv*_ -specific primers (dGFPuvF2/dGFPuvR2) to detect shuttle vector. Lane designations: 1: no template control; 2: pRMΔ2 passage 1 transformant; 3. pRMΔ2 passage 3 transformant; 4. pRMΔ1 passage 3 transformant; 5. pRMΔ3 passage 3 transformant; 6. WT *R. monacensis*; 7: *R. amblyommatis* pRMΔ2 transformant.

### Copy number ratios of native pRM and pRM shuttle vectors in transformed *R. monacensis*

Quantitative PCR (qPCR) estimation of the relative ratio of the single copy pRM-encoded resolvase RM_p5 (qRM_p5F/ qRM_p5R primer set, Table 2) and chromosome-encoded *gltA* genes (29) indicated 1.5 copies of pRM for each chromosome copy in WT *R. monacensis*, which decreased to a ratio of 0.5-0.8 in pRMΔ1, 2 and 3 transformed-*R. monacensis* (Fig. 5). This ratio was mirrored by the 0.5-0.9 ratio of shuttle vector (qGFPuvF/qGFPuvR primers from shuttle vector-encoded *gfp*_*uv*_, Table 2) to chromosome in the transformant rickettsiae (Fig.5). There were no further significant changes in copy number ratios during serial passage of the transformant rickettsia (up to 20 passages in the case of pRMΔ2)(data not shown).

**FIG 5.**
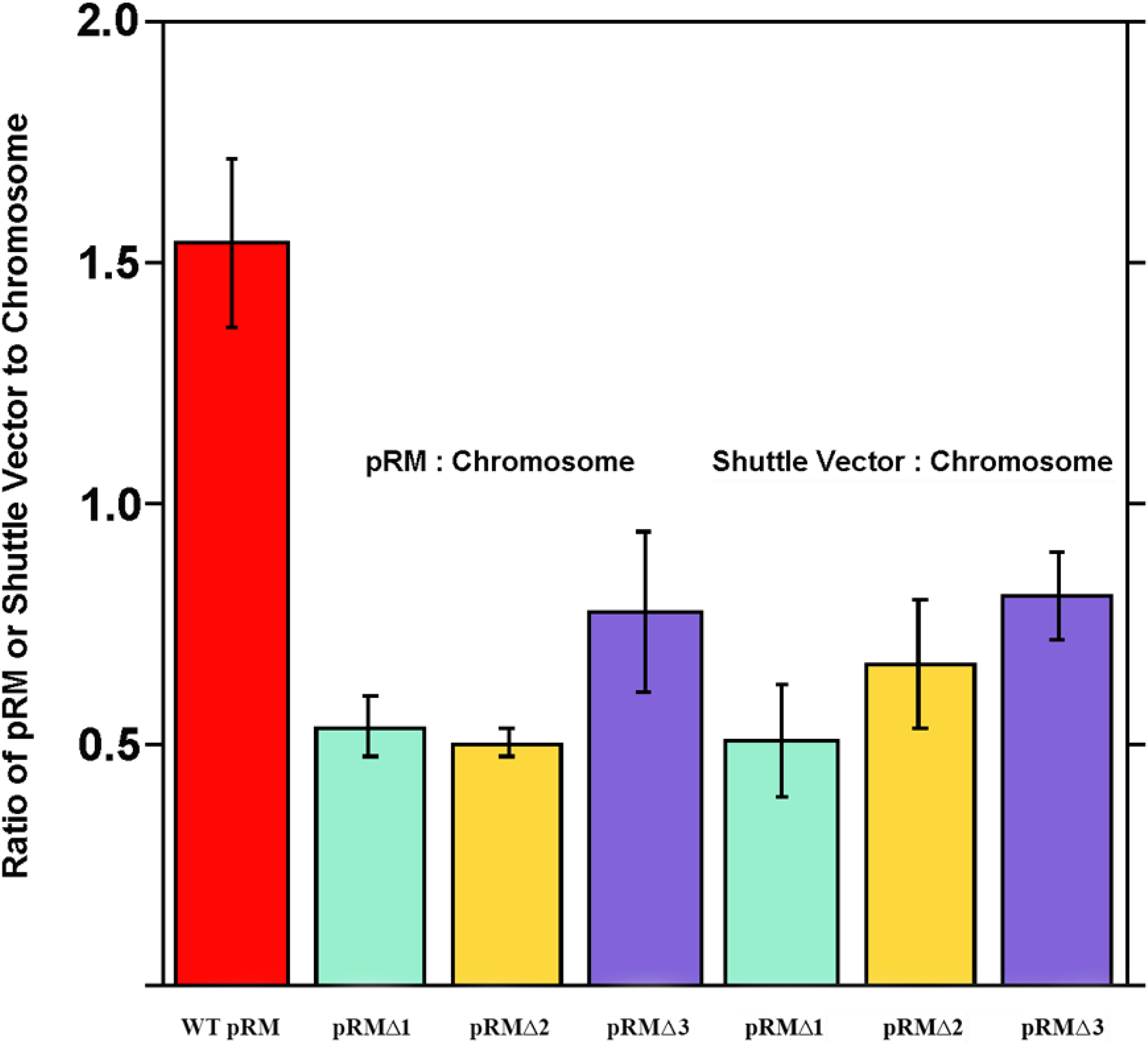
**Plasmid to chromosome copy number** ratios in transformed *R. monacensis*. Native plasmid (pRM), shuttle vector and chromosomal copy numbers were quantified by qPCR of *RM_p5, gfp*_*uv*_ and *gltA* respectively. Passage numbers (p) used for this assay were as follows: WT p 17 (all 3 replications); pRMΔ2 p 10 (all 3 replications); pRMΔ1 one replication with p 3 and 2 replications with p 9; pRMΔ3 one replication with p 3 and 2 replications with p 7.

### Presence of shuttle vector and native plasmid complexes in pRM-transformed *R. monacensis*

To confirm the PCR-indicated presence of native pRM in pRMΔ shuttle vector-transformed *R. monacensis*, undigested genomic DNA from WT and pRMΔ1, 2 and 3 transformants was separated by pulsed-field gel electrophoresis (Fig. 6) and transferred onto zeta probe membranes. Nicked linear, circular and multimeric forms of pRM were present in the WT *R. monacensis* lanes (Fig. 6A and C; bands with white asterisks). Presence of the shuttle vector in *R. monacensis* transformed pRMΔ1, pRMΔ2 and pRMΔ3 was demonstrated by hybridization with digoxigenin-labeled *gfp*_*uv*_ probe (absent in the WT) (Fig. 6B). The *gfp*_*uv*_ probe localized to bands in the ∼40 kbp region (arrowhead) rather than at 10-12 kbp (the size of the shuttle vectors), indicating that there are no detectable monomeric pRMΔ1, pRMΔ2, or pRMΔ3 in transformant populations.

**FIG 6.**
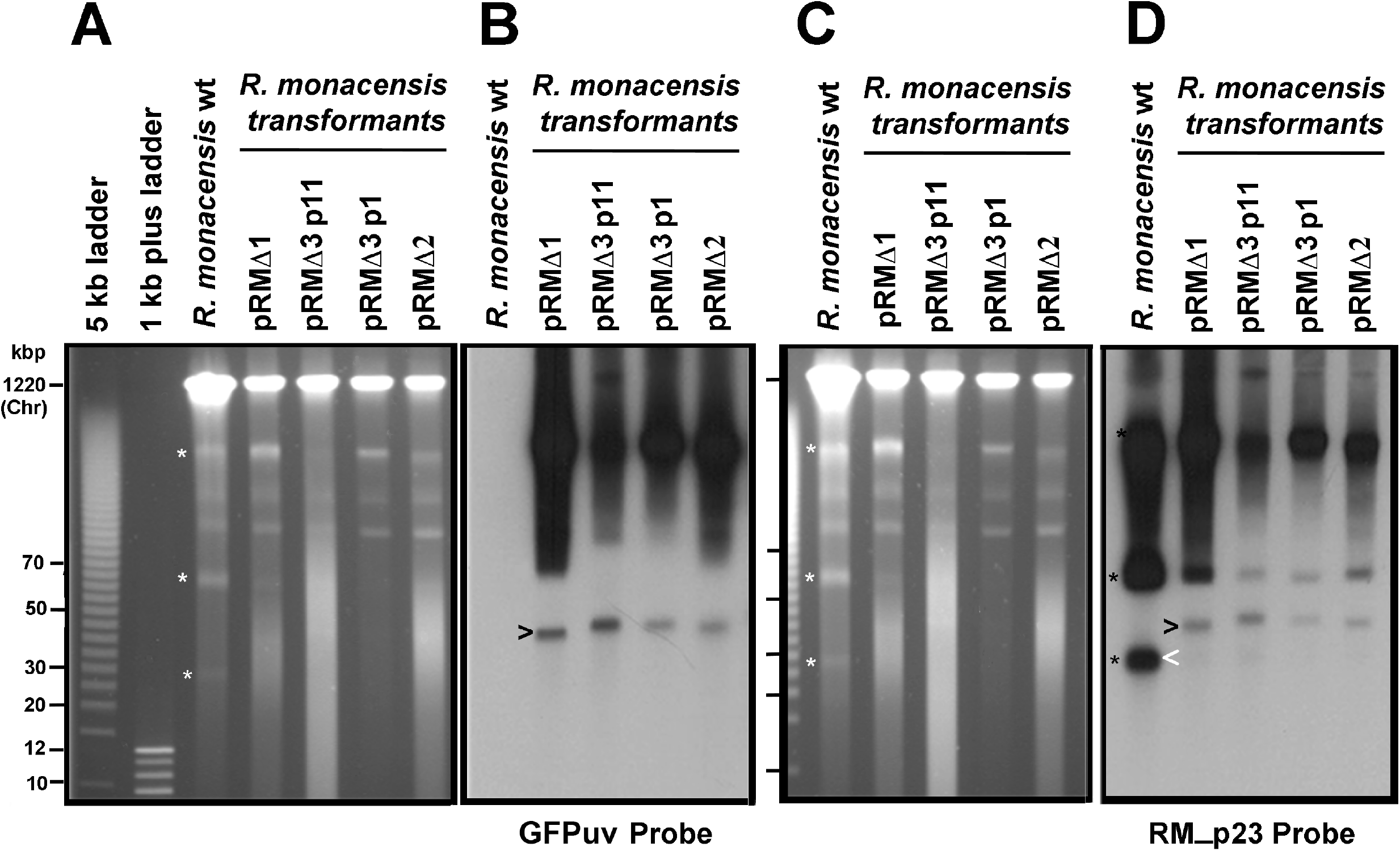
Pulsed-field gel electrophoresis and Southern blot analysis of *R. monacensis* pRM shuttle vectors transformants. A) PFGE gel and B) Southern blot analysis of panel A hybridized with digoxigenin-labeled *gfp*_*uv*_ probe showing the presence of shuttle vectors (black arrowhead indicates nicked linear forms). C) PFGE gel and D) Southern blot analysis of panel C hybridized with digoxigenin-labeled RM_p23 probe showing the presence of native pRM in the WT *R. monacensis* (white arrowhead indicates nicked linear form) and a larger form (black arrowhead) in each of the transformants that matches the size of the products that hybridized with the *gfp*_*uv*_ probe (panel B). White asterisks in panels A & C mark the 3 main forms of native pRM on the PFGE gels, while the black asterisks show corresponding hybridization of said forms with RM_p23 probe in panel D. p1 and p11 refer to the passage number of pRMΔ3.

The PFGE gel shown in Figure 6C was transferred to a membrane and hybridized with a probe containing digoxigenin-labeled RM_p23 (13), a gene present on native pRM but absent on the shuttle vectors (Fig. 6D). In the WT lane, the probe hybridized to the 20-30 kbp linear nicked and multimeric forms (black asterisks) of pRM. In the transformant lanes, the RM_p23 probe hybridized in the ∼40 kbp region (Fig. 6D, arrowhead) at the same relative positions as the *gfp*_*uv*_ probe (Fig. 6B). The strong hybridization of the RM_p23 probe at the 40 kbp position in transformant rickettsiae lanes versus weak hybridization at the lowest position (lowest black asterisk panel D) indicates likely presence of native pRM that is predominantly complexed with shuttle vector, as transformants lacking pRM would not hybridize with RM_p23.

## DISCUSSION

Although plasmids have been extensively studied in other bacteria, the discovery of plasmids in rickettsiae is fairly recent (1, 2, 13), and their biological function is largely unexplored. Genomic analysis has shown that rickettsiae have undergone reductive evolution (30-33) resulting in characteristics such as AT enrichment, high conservation of genome sequences among species, higher levels of virulence, and variable presence/numbers of plasmids (3). Rickettsial plasmids have likewise undergone reductive evolution and mirror rickettsial genomes in relative size and GC content (34). It has been proposed that a rickettsial ancestor supported a plasmid system which was lost in some species due to their unique obligate intracellular lifecycle (3, 34), but other species presumably retained plasmids due to an advantage conferred by their presence. Plasmids are known agents of horizontal gene transfer, facilitating host adaptation/virulence, antibiotic and stress resistance, and genetic plasticity (1, 34, 35, 36). Though origins and functions of many rickettsial plasmid gene sequences have been inferred from similarities to those of other bacteria, the role of plasmids and their interactions in rickettsiae remains to be elucidated. Real-time PCR and whole genome sequencing showed that these are low copy number plasmids (2), while creation of shuttle vectors from pRAM18 and pRAM32 and their subsequent transformation into rickettsiae confirmed that *parA* and *dnaA* were essential for plasmid replication and maintenance (12). However, the mechanism by which these genes function is yet to be identified.

We have developed two new *Rickettsia* plasmid shuttle vectors, pRMΔ2 andΔ3, that can be used to transform plasmid-free *Rickettsia* spp. as well as those carrying native plasmids. In conjunction with pRMΔ1 they collectively contained varying regions of pRM surrounding the *dnaA* and *parA* genes. Analysis of their relative efficacy in transformed rickettsiae allowed us to identify plasmid sequences important for the replication and maintenance of the shuttle vectors in rickettsiae. Specifically, the inability of *R. parkeri, R. bellii, R. montanensis* (plasmid-free), and *R. amblyommatis* strain AaR/SC (carrying 3 plasmids) to transform with shuttle vector pRMΔ1 (Figure 1C, containing RM_p16 through RM_p19 and the 3’ end of RM_p20) while transforming easily with shuttle vectors pRMΔ2 andΔ3 (containing RM_p16 through RM_p20 and pRM21 respectively) indicated that RM_p20 or its immediate upstream sequence was required for shuttle vector replication in rickettsiae. These data support the prediction (37) that the region of pRM containing RM_p17 to RM_p20 forms an operon. It is possible that RM_p19 and 20 are not themselves required because absence of a single promoter upstream of RM_p20, as predicted by BPROM (25), would prevent expression of RM_p18 (*parA*) in pRMΔ1. Without expression of the ParA chromosomal stability protein the shuttle vector would not be maintained, consistent with the observed phenotype. In contrast, *R. monacensis* was transformed with pRMΔ1 when the other species were not, likely due to the presence of native pRM providing the necessary ParA for maintenance of the shuttle vector.

Partition systems usually include three features: a site that acts like a centromere, a centromere-binding protein (CBP) (usually encoded by *parB*) and an NTPase (*parA*) (38). As none of the genes in the pRM operon have similarity to known *parB* genes, the pRM partitioning system could work in one of several ways. The partitioning system could rely on the chromosomal *parB* to work or represent a new type of system, as in R388 which functions with a single protein and a centromere site (38). Alternatively, the pRM hypothetical genes could act in the capacity of *parB* as CPBs are not necessarily significantly similar in sequence but are typically dimers of HTH or ribbon helix-helix DNA-binding proteins (38). Both RM_p19 and 20 have HTH_XRE domains and might function as CPBs.

The pRM operon containing *parA* appears to be unique amongst known Rickettsial plasmids. A BLASTN of the genes RM_p17 and 20 showed no rickettsial homology, while RM_p19 only had homology with the Rickettsial endosymbiont of *Ixodes pacificus* plasmid (pREIP). Interestingly, translation of the nucleotide sequence for RM_p19 and 20 and subsequent BLASTP of the amino acid sequence shows a low level of similarity to pREIP (36% ID, 59% positive for RM_p19 and 40% ID, 60% positive for RM_p20) and to Rickettsial endosymbionts of a variety of arthropods, including beetles such as *Platyusa sonomae* and *Bembidion* nr Transversale, bedbugs (*Cimex lectularis*) and midges (*Culicoides impunctatus*). These *Torix*-clade *Rickettsia* spp. (39) have 2 or 3 different regions of similarity for both RM_p19 and 20 and have IDs ranging from 30-40% with 53 to 62% positives.

The ParA protein encoded on *R. monacensis* pRM is sufficiently distinct from those of pRAM18, 23 and 32 (the native plasmids of *R. amblyommatis*) that it should support uptake and maintenance of the pRM shuttle vector in *R. amblyommatis*. In their comparison of rickettsial plasmid genes, El Karkouri et al. (3) showed that pRM *parA* was distinct from the *parA* of all other sequenced rickettsial plasmids. A BLASTP search of the ParA protein from pRM identified only one rickettsial homolog (*Rickettsia asembonensis* with 46% identity), while its closest match was from a Mycoplasmataceae bacterium, with 53% identity. Lack of interactions between non-homologous ParA proteins may explain why *R. amblyommatis* strain AaR/SC was transformed with pRM shuttle vectors despite its three native plasmids and their potential for causing plasmid incompatibility. On the other hand, the successful transformation of *R. monacensis* with its own pRM shuttle vectors was unexpected. It is possible that the presence of identical ParAs from native plasmids and shuttle vectors supports mutual plasmid maintenance in rickettsiae rather than promoting incompatibility. These results also highlight the fact that plasmid transformation of different *Rickettsia* species is by no means a routine activity with universally predictable results. For example, the construct pRAM18/Rif/GFPuv containing full-length pRAM18 successfully transformed *R. bellii* and *R. parkeri* (12), but for unknown reasons, shuttle vectors that incorporated full-length pRM instead of pRAM18 (data not shown) were unable to transform *R. montanensis, R. monacensis, R. peacockii, R. parkeri* or 3 strains of *R. amblyommatis*. Thus, there is still much to learn about the role of these rickettsial plasmids in the functioning of rickettsiae, and the mechanisms by which they operate.

The studies reported here give us a better understanding of the mechanisms of rickettsial plasmid maintenance in diverse rickettsial species. We explored the basis for the ability of pRM based shuttle vectors to transform *R. monacensis*. Our PCR results detected the presence of shuttle vector in transformants which continued to persist through serial transferring. They also indicated a decrease in the ratio of pRM to the chromosome suggesting that transformants harbored an average of two or more chromosome copies per cell, or that plasmids were cured from some of the rickettsiae. Nevertheless, data from Figure 2 suggests that almost 100% of the rickettsiae contained the shuttle vector. Furthermore, the results showed that the use of homologous rickettsial *parA* regions leads to the formation and maintenance of complexes between shuttle vectors and native plasmids, suggesting possible defects in partitioning of plasmids carrying the same *parA* genes. The surprising ability of *R. monacensis* to be transformed by shuttle vectors containing the *parA* gene from its native plasmid emphasizes the need for further study of plasmid maintenance and incompatibility in rickettsiae and further studies are needed to elucidate the molecular basis for the apparent linkage of shuttle vectors with pRM present in *R. monacensis*.

## MATERIALS AND METHODS

### Bacterial strains and plasmids

*Rickettsia monacensis* strain IrR/Munich^T^ (40) WT was used at passages 10-70 after initial isolation from a tick. It was propagated in *Ixodes scapularis* cells, line ISE6 as described (16). *R. amblyommatis* (strain AaR/SC), *R. bellii* (strain RML 369-C), *R. parkeri* (strain Tate’s Hell), and *R. montanensis* (strain M5/6) were propagated in the same manner as *R. monacensis*.

The *R. monacensis* plasmid, pRM, was cloned in *E. coli* as previously described (37). Briefly, electroporation of *R. monacensis* with the pMOD658 transposon yielded the Rmona658B transformant in which the pMOD658 transposon encoding chloramphenicol acetyltransferase (CAT) and a *gfp*_*uv*_ fluorescent marker was inserted into pRM (37). The mutated pRM was cloned in its entirety by chloramphenicol marker rescue of Big Easy TSA™ *E. coli* cells (Lucigen, Middleton, WI) electroporated with Rmona658B genomic DNA that had been linearized by digestion with *Sma*I and ligated into the linear vector pJAZZ (Lucigen). The resulting construct was termed pJAZZ[pRM658B] (Fig. 1A).

The following abbreviations are used to designate plasmids and their derivatives: “S” indicates spectinomycin/streptomycin resistance conferred by the aminoglycoside adenyltransferase gene, *aadA*, driven by a rickettsial *ompA* promoter; “G” indicates green fluorescent protein encoded by *gfp*_*uv*_ under regulation of the rickettsial *ompA* gene promoter; “K” indicates kanamycin resistance from the pET-28a (+) vector (Novagen, EMD Millipore, Bedford, MA) adapted as described below. Specific nucleotide spans from pRM are from Acc. EF564599 (37). Multiple cloning sites (MCS) are designated [MCS]. Promoters are designated *p*.

### Construction of pRM-based shuttle vectors: preparation of selection reporter cassette

We constructed a cassette into which pRM fragments could be cloned. The SGK selection reporter cassette (Fig. S1C) was derived from the previously constructed shuttle vector pRAM18dRGA[MCS] (12) (Fig. S1A) and contained genes needed for replication and antibiotic selection in *E. coli* as well as reporter and antibiotic selection genes for use in rickettsiae (Fig. S1C). Replacement of pGEM with a 3.159 kbp DraIII/PshA fragment of the pET-28a (+) vector was prompted by previous experiments indicating that some genes cloned into the pRAM18 shuttle vector MCS were less stable in pGEM than pET. Because spectinomycin and streptomycin are water soluble and not used to treat rickettsioses but rifampin may be, we replaced the *rpsLp-arr-2*_*Rp*_ (RIF)/*ompAp*-*gfp*_*uv*_ cassette with an *ompAp-aadA*/*ompAp*-*gfp*_*uv*_ cassette (Fig. S2B).

### Construction of pRM-based shuttle vectors

Sub-cloning pRM: To identify which region(s) of pRM yielded the most effective shuttle vector for rickettsial transformation we cloned specific fragments of pRM to create a family of deletion constructs, pRMΔ1, pRMΔ2 and pRMΔ3. To construct pRMΔ1 and pRMΔ3, pJAZZ[pRM658B] was digested with BamHI/PacI or BamHI /PflMI (Fig. 1A and B) and the desired 4671 bp and 7177 bp fragments containing either RM_p16 to 20 (pRM bp 14561-19231) or RM_p16 through 21 (pRM bp 14561-21646) were gel-purified (Zymoclean ™Gel DNA Recovery Kit, Zymo Research, Irvine, CA). The fragment ends were blunted (DNATerminator®) and ligated into the blunt and dephosphorylated selection reporter cassette (SGK), creating 10368bp pRMΔ1 and 12797bp pRMΔ3, respectively (Fig. 1D). To construct pRMΔ2, Q5 DNA polymerase (New England Biolabs) was used to PCR-amplify RM_p16 through RM-p20 and into the 5’ end of RM_p21 (pRM bp 14512 through 19585) with primers RM_p16 FOR NheI/ RM_p21Rev NheI (5’-TAT TGC TAG CCG TAA GGA ACA GTT GGT GAG -3’ and 5’-ATA TGC TAG CGT TAA TAT GCC TCG GGC TAC -3’). The PCR product with NheI sites at both ends (Fig. 1C) was incubated with taq polymerase to create A’ overhangs and cloned into pCR4 with the TOPO TA Cloning® Kit (Invitrogen, Carlsbad, CA). Clones were sequenced to verify that they contained the correct pRM fragment which was then recovered by restriction digest with NheI and ligated it into dephosphorylated, NheI-digested SGK, to yield the 10881 bp shuttle vector pRMΔ2 (Fig. 1D).

To predict the location of rickettsial promoters in the pRM fragments cloned into the shuttle vectors, we used BPROM (25) (*http://www.softberry.com/berry.phtml?topic=bprom&group=programs&subgroup=gfindb*).

### Preparation of shuttle vector plasmid DNA

Endotoxin-free maxi preps (Qiagen, Valencia, CA) were prepared for all pRMΔ shuttle vectors as per manufacturer’s recommendations, for use in transforming rickettsiae. Integrity and orientation of inserted genes was re-confirmed by sequencing fragment junctions and by restriction digest analysis. (See Table 3 for sequencing primers).

**TABLE 3.**
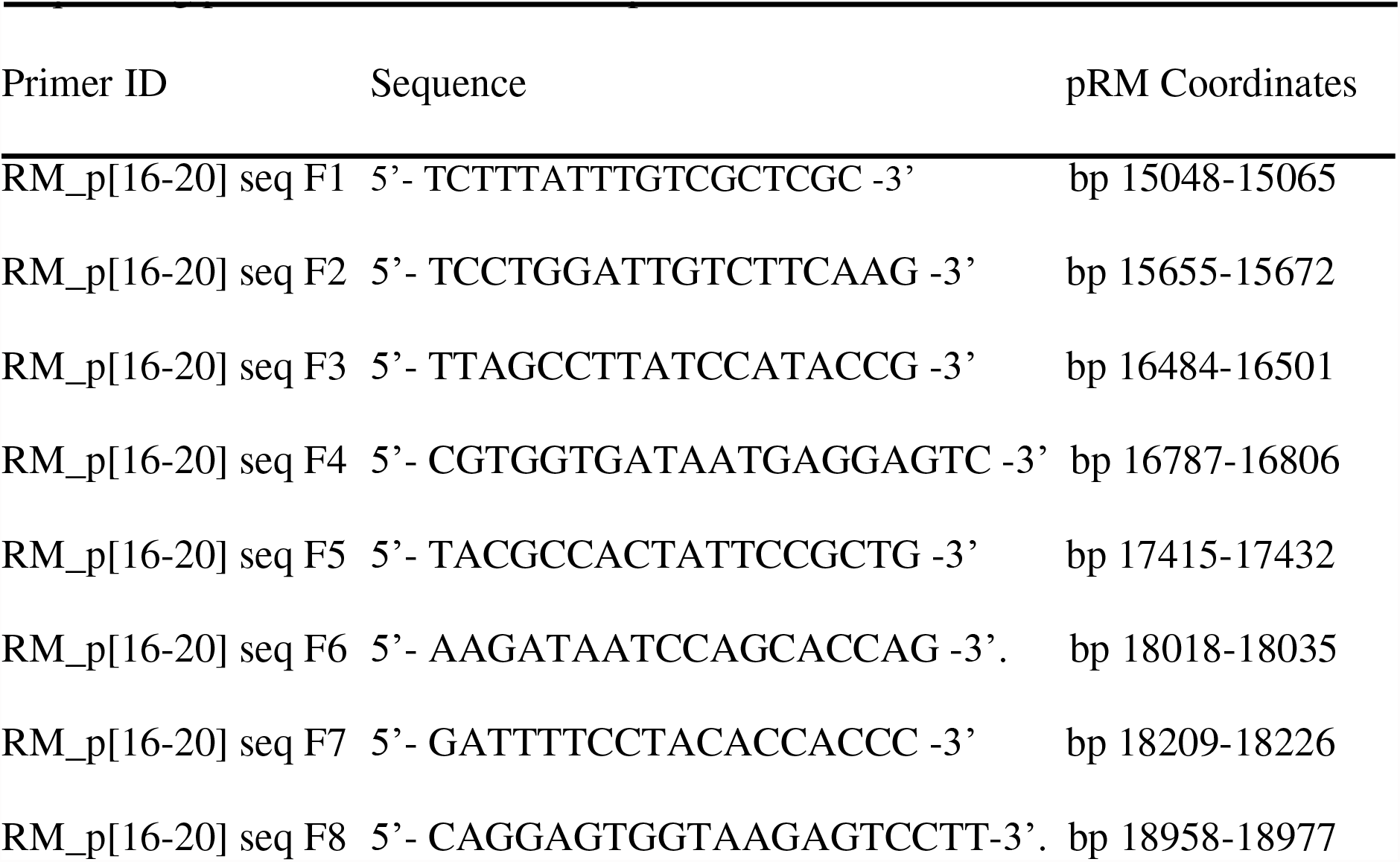
Sequencing primers for shuttle vector pRMΔ2

### Transformation of rickettsiae

Rickettsiae were purified, and electroporated as described (16). Rickettsial transformants were selected using growth medium containing spectinomycin and streptomycin each at a final concentration of 100 μg/ml. Cultures were monitored for expression of the GFP_uv_ reporter on a weekly basis using an inverted microscope (Nikon Diaphot TMD with Y-FL-epifluorescence attachment and a sapphire GFP 31043 filter), or by examining wet mounts on an upright Nikon Eclipse E400 microscope with a B-2E/C FITC filter (Nikon, Melville, NY).

### Growth rate analysis of *R. monacensis* transformed with pRMΔ2

Multiwell plates of ISE6 cells were inoculated with *R. monacensis* WT at a 1:25 dilution and sampled at selected times (2.8, 48.5, 72.8, 96.4, 119.7, 143.7, 168.9 and 191.4 hours post-inoculation [hpi]). Two plates of *R. monacensis* WT diluted 1:50 were sampled at 0, 22.8, 47.2, 70.3, 94.5, 119.8, 137.7, and 167 hpi. One plate of ISE6 cells inoculated with the *R. monacensis* pRMΔ2 transformant diluted 1:25 was sampled at 4, 49.5, 97.2, 145.3, 215.5, 264.2, 311.5 and 359.5 hpi. Two additional plates seeded with the same transformant were diluted 1:10 and 1:25 and sampled at 3.3, 72.1, 98.5, 121.2, 145, 167.2, 217.8 & 265.4 hpi. Growth rates were based on qPCR-determined levels of chromosomal single copy *gltA* (29) from the average of 3 growth curve analyses.

### Evaluation of GFP expression in WT and transformed *R. monacensis* using confocal microscopy

Cell-free WT and transformed *R. monacensis* were resuspended in complete medium and incubated with NucBlue™ Live Cell Stain ReadyProbes™ reagent (1: 50 dilution) (Thermo Fisher Scientific) in the dark for 30 min at room temperature. 50 μl aliquots of cell-free *R. monacensis* were deposited onto microscope slides (Cytospin centrifuge, Thermo Fisher) at 200 rpm for 3 minutes. The slides were mounted with 3 μl 1X PBS and imaged on an Olympus BX61 DSU confocal microscope with a 60X objective via a double-wavelength filter [4, 6-diamidino-2-phenylindole (DAPI): excitation at 365 nm and emission at 480 nm; FITC: excitation at 495 nm and emission at 519 nm]. Co-localization of fluorescence from *gfp*_*uv*_ (transformed *R. monacensis*) and NucBlue™ (rickettsial DNA) was analyzed by determining signal overlap for each of three random fields of view using Image Fiji (JaCoP plugin and Co-localization Threshold plugin), Pearson’s coefficient (PCC) and calculation of Manders’ co-localization coefficients (MCC) (26, 27).

### Pulsed-field gel electrophoresis and Southern blot analyses

Concentrations of cell-free rickettsiae were estimated using OD600 values. Approximately 1.0-2.0 × 10^9^ bacteria were embedded in 1% Incert agarose (BMA, Rockland, MD) and rickettsial DNA was released as previously described except incubations were performed at 50ºC (37). Agarose plugs were equilibrated with 0.5X TBE, and ½ of each plug was inserted into a well of a 1% LE agarose gel (Beckman Instruments, Inc., Palo Alto, CA) and electrophoresed on a CHEF Mapper® XA Pulsed-Field Gel Electrophoresis System (BioRad, Hercules, CA) as previously described except the setting for “molecular weight low” was 5kbp (12). Depurinated gels were transferred onto Zeta Probe GT genomic membranes (Bio-Rad) and hybridized (50ºC), washed, labeled, detected, stripped and re-hybridized as previously described (12, 13, 28, 37). DNA probes used were specific for genes encoding GFP_uv_ to detect shuttle vector (28), RM_p23 to detect native pRM plasmid (12), and recombinase (invertase), Hsp2 and helicase (RecD) to detect *R. amblyommatis* strain AaR/SC plasmids pRAM18, 23 and 32 respectively (12).

### PCR and sequence confirmation of rickettsial species and copy number estimates of native pRM and pRM shuttle vectors in transformants

To confirm species identities, portions of the *ompA* gene of spotted fever group rickettsiae were amplified using primers 190-70/190-602 (41) and rickettsial genomic DNA as template (42). PCR products were purified with the DNA Clean and Concentrator kit (Zymo Research) as per manufacturer’s protocol, for Sanger sequencing on an ABI 3730 Excel automated sequencer (University of Minnesota Genomics Center). The sequences were compared to rickettsial *ompA* sequences in GenBank using BLASTN® (NCBI, NIH).

Three sets of specific primer pairs (Table 2) were designed for PCR amplification of native pRM while the dGFPuvF2/R2 primer pair (Table 2) was used to amplify a product specific to pRM shuttle vectors in rickettsiae. PCR reactions used 100 ng of template, 1 mM mixed dNTPs, 0.5μM of each primer, and 1.25 units of GoTaq polymerase units in 50 μl final 1X PCR buffer (Promega, Madison, WI), and were run in a Robocycler (Stratagene, La Jolla, CA) as follows: 1 cycle 95ºC for 3 min, 40 cycles 95ºC for 15 s, 51ºC for 30 s, 72ºC for 1 min, and one final 7 min cycle at 72ºC.

Real-time quantitative PCR (qPCR) was used to estimate relative copy number ratios (43, 44) of single copy genes specific to the *R. monacensis* chromosome (*gltA* encoding citrate synthase), the native pRM plasmid (RM_p5 locus for transposon resolvase), and the pRM-derived shuttle vectors (GFP_uv_) using three primer pairs (Table 2). With the exception of a 56ºC annealing temperature, reactions and copy number estimates were executed as previously described (2).

## ACKNOWLEDGEMENTS

This research was supported by grants (R01 AI049424 and R01 A1081690) to UGM from the USA National Institutes of Health. We thank Roderick Felsheim for technical advice and assistance.

